# Diversification of the recombinant anti-kinesin monoclonal antibody H2

**DOI:** 10.1101/2022.12.22.521561

**Authors:** Shinsuke Niwa, Kyoko Chiba

**Affiliations:** Frontier Research Institute for Interdisciplinary Sciences (FRIS), Tohoku University, Aramaki-Aoba 6-3, Aoba-Ku, Sendai, Miyagi 980-0845, Japan

**Author notes:** Corresponding authors: Shinsuke Niwa, Frontier Research Institute for Interdisciplinary Sciences (FRIS), Tohoku University, Aramaki-Aoba 6-3, Aoba-Ku, Sendai, Miyagi 980-0845, Japan, Kyoko Chiba, Frontier Research Institute for Interdisciplinary Sciences (FRIS), Tohoku University, Aramaki-Aoba 6-3, Aoba-Ku, Sendai, Miyagi 980-0845, Japan. These authors equally contribute to this work.

## Abstract

Kinesin-1, a motor protein composed of the kinesin heavy chain (KHC) and the kinesin light chain (KLC), is fundamental to cellular morphogenesis and function. A monoclonal antibody (mAb) called H2 recognizes the KHC in a broad range of species and is one of the most widely used mAbs in cytoskeletal motor research. Here, we generated vectors that expressed recombinant H2 in mammalian cells. We demonstrated that the recombinant H2 performed as well as the hybridoma-derived H2 in western blotting and immunofluorescence assays. The recombinant H2 could detect all three human KHC isotypes (KIF5A, KIF5B, and KIF5C) and amyotrophic lateral sclerosis (ALS)-associated KIF5A aggregates in the cell. Immunofluorescence microscopy showed that the single chain variable fragment (scFv) derived from the H2 mAb could specifically recognize KHCs in cells. In addition, we developed a chickenized anti-KHC scFv(H2), which broadens the application of H2 in immunofluorescence microscopy. Collectively, our findings validate recombinant H2 as useful for studying the function of KHCs.

## Introduction

Kinesin superfamily proteins (KIFs) are microtubule-dependent molecular motors (Hirokawa *et al*., 2009). KIFs have diverse roles in the cell, such as determining the localization of organelles, exerting forces during cell division, and stabilizing and depolymerizing microtubules (Okada *et al*., 1995; Evans *et al*., 2006; Pan *et al*., 2006; Niwa *et al*., 2008; Scholey, 2008; Zhou *et al*., 2009; Niwa *et al*., 2012). Kinesin-1, the first KIF to be identified, was originally attributed the function of a molecular motor for axonal transport (Brady, 1985; Vale *et al*., 1985; Aizawa *et al*., 1992). Previous studies have shown that kinesin-1 is evolutionarily conserved and is involved in the transport of various cellular components including the nucleus, mitochondria, lysosomes, neurofilaments, and RNA granules in the cell (Vale *et al*., 1985; Tanaka *et al*., 1998; Xia *et al*., 2003; Kanai *et al*., 2004; Uchida *et al*., 2009). Recent studies have shown that the intracellular transport of some viruses by kinesin-1 is an important step in viral infection and maturation (Morgan *et al*., 2010; Pegg *et al*., 2021; Xu *et al*., 2023).

Kinesin-1 is a tetramer composed of two subunits of kinesin heavy chain (KHC) and two subunits of kinesin light chain (KLC) (Bloom *et al*., 1988). *Drosophila melanogaster* and *Caenorhabditis elegans* have one KHC gene each (Yang *et al*., 1989; Patel *et al*., 1993), while mammals have three KHC genes: *KIF5A, KIF5B*, and *KIF5C. KIF5A* and *KIF5C* are neuron-specific isotypes of KHC (Kanai *et al*., 2000; Xia *et al*., 2003), whereas *KIF5B* is a ubiquitous isotype that is expressed in a broad range of cell types (Tanaka *et al*., 1998). We have recently shown that KIF5A, KIF5B, and KIF5C have different motilities and oligomerization tendencies (Chiba *et al*., 2022). KIF5B and KIF5C have shorter run lengths and lower microtubule binding rates than KIF5A. In addition, only KIF5A shows a strong propensity to form oligomers. It has been suggested that this property of KIF5A is related to the pathogenesis of amyotrophic lateral sclerosis (ALS), which is caused by *KIF5A* mutations (Baron *et al*., 2022; Nakano *et al*., 2022; Pant *et al*., 2022). The mutation of *KIF5B* and *KIF5C* is also associated with genetic disorders. The mutation of *KIF5C* causes neurodevelopmental disorders (Poirier *et al*., 2013; Michels *et al*., 2017; Duquesne *et al*., 2020), while the mutation of *KIF5B* is responsible for a broader spectrum of diseases including skeletal dysplasia, dilated cardiomyopathy and skeletal myopathy (Flex *et al*., 2022; Itai *et al*., 2022). Therefore, researchers in a wide range of disciplines (including basic cell biology, neuroscience, virology, and medical science) are currently studying KHCs and kinesin-1.

Monoclonal antibodies (mAbs) are useful tools in biology and medicine. mAbs that recognize KHCs were generated using purified kinesin-1 as an antigen (Pfister *et al*., 1989). An anti-KHC mAb called H2 is the most commonly used mAb in microtubule-dependent motor research. This is because H2 recognizes KHCs from a broad range of species. In addition, H2 can be used in wide variety of applications, such as immunofluorescence microscopy, western blotting, functional kinesin-1 blocking, and immuno-electron microscopy (Hirokawa *et al*., 1989; Brady *et al*., 1990). While purified H2 can be purchased from commercial vendors, it is very expensive. An H2 hybridoma is also available upon request (Pfister *et al*., 1989); however, transporting living or frozen cells is not always easy, especially when requests are made from outside the USA. In recent years, it has been possible to generate recombinant mAbs, which overcome these problems(Andrews *et al*., 2019). When an immunoglobulin (Ig)G cDNA sequence of an antibody is determined, recombinant antibodies and their derivatives can be produced indefinitely. Researchers can produce large-scale yields of recombinant antibodies by themselves using cost-effective expression systems such as HEK293 cells. Moreover, plasmids are much more easily preserved and distributed than hybridoma cells. Here, we determined the sequences of IgG cDNA from H1 and H2 hybridomas. Using the H2 IgG cDNA sequence, an expression plasmid for recombinant H2 antibody was generated. Western blotting and immunofluorescence were used to demonstrated that the recombinant H2 antibody worked as effectively as the antibody obtained from the H2 hybridoma. An additional advantage of recombinant antibodies is the potential for their manipulation to create new versatile reagents. Thus, using the primary amino acid sequence of H2 IgG, we generated a single chain variable fragment (scFv) of H2 (scFv[H2]). By fusing scFV(H2) and the Fc region of chicken IgG, we were further able to change the isotype of H2.

## Results and Discussion

### Cloning of H1 and H2 variable regions and construction of recombinant H2 expression plasmid

IgG cDNA was obtained by 5’ RACE as previously described (Meyer *et al*., 2019). The Kappa chain but not the lambda chain fragment was obtained from both H1 and H2 hybridomas. This is consistent with the composition of the H1 and H2 IgGs previously reported (Pfister *et al*., 1989). The variable region sequences were determined by Sanger sequencing (Fig 1A and 1B).

**Figure 1.**
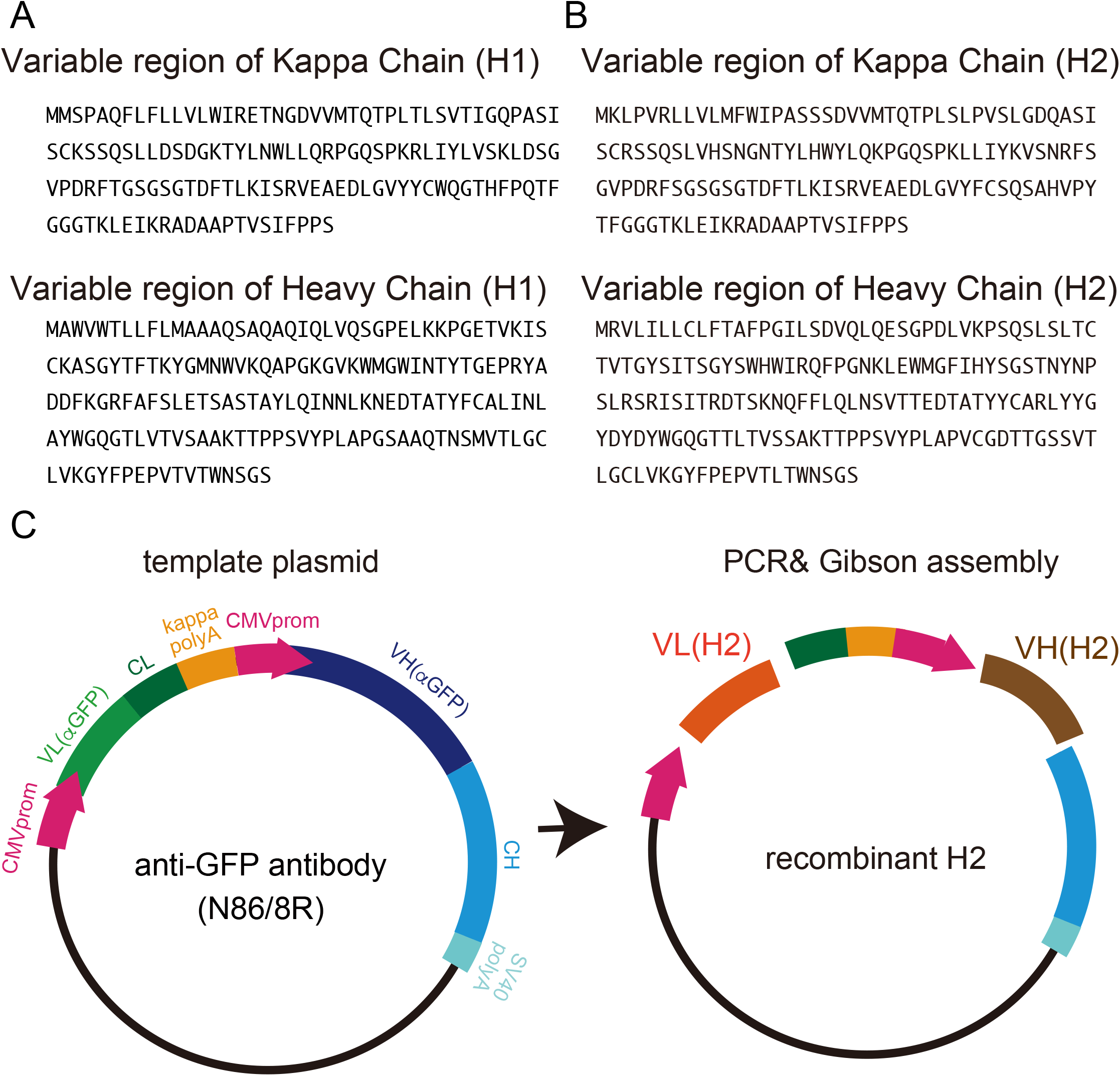
Variable regions of anti-kinesin-1 antibodies and the H2 IgG expression plasmid. (A and B) Amino acid sequences of the variable regions of the IgG light and heavy chains from H1(A) and H2 (B). (C) Schematic drawing showing the production of the recombinant H2 IgG expression plasmid. The recombinant H2 expression plasmid was generated using the anti-GFP antibody expression plasmid as a template.

We then generated a plasmid to express recombinant H2. To this end, we swapped the variable regions of the expression vector for the recombinant N86 antibody targeting the green fluorescent protein (GFP) (Andrews *et al*., 2019) (Fig 1C).

### Use of recombinant H2

To express recombinant H2, the expression vector was introduced into 293FT cells. Three days later, the culture medium was obtained. The experiments described in this report were performed using the cultured medium because we found that purification was not required for western blotting or immunofluorescence microscopy. We first performed western blotting. Purified human KIF5A, KIF5B, and KIF5C expression was analyzed using the recombinant H2. We found that although the recombinant H2 recognized all three human KHCs, the KIF5B-specific signal was much weaker than those of KIF5A and KIF5C (Fig 2A). This result is consistent with that of a previous study performed with the original H2 mAb (Kanai *et al*., 2000), suggesting that the specificities of the original and recombinant H2 mAbs were similar.

**Figure 2.**
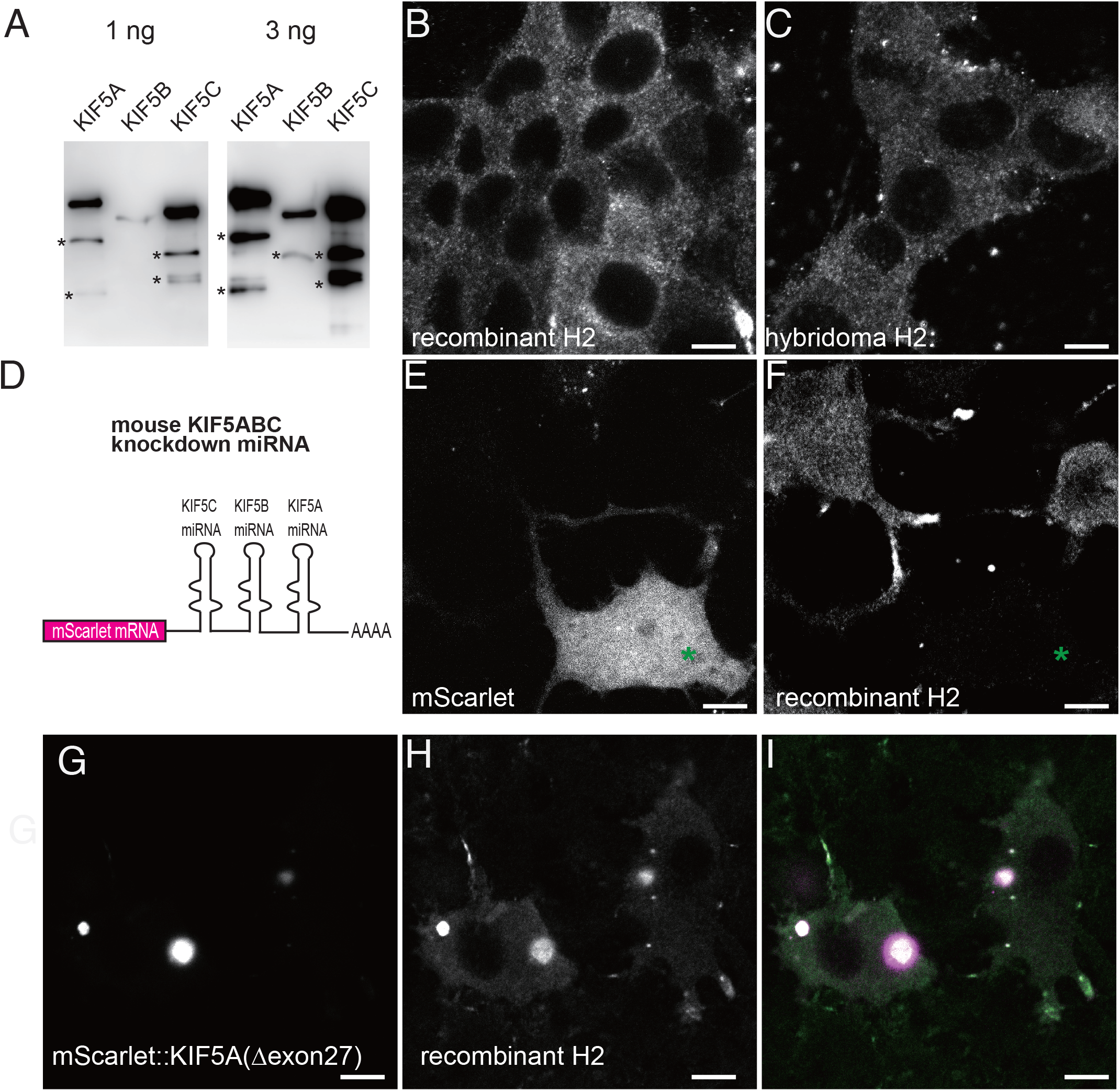
Characterization of recombinant H2. (A) Western blotting using 293FT-cell-derived recombinant H2. 1 ng (left) and 3 ng (right) of purified human KIF5A, KIF5B, and KIF5B were separated by SDS-PAGE and western blotting was performed. Asterisks indicate degraded proteins recognized by recombinant H2. (B and C) Comparison of recombinant H2 (B) and hybridoma H2 (C) in immunofluorescence microscopy images of stained CAD cells; bars = 10 μm. (D–F) The function of the recombinant H2 antibody was validated by the simultaneous knockdown of three KHC isotypes. (D) Schematic drawing showing mScarlet mRNA fused with miRNAs that simultaneously knocked down three KHC isotypes, *KIF5A, KIF5B*, and *KIF5C*. (E and F) CAD cells were transfected with the KHC knockdown vector and stained with recombinant H2; the mScarlet fluorescence signal (E) and the recombinant H2 fluorescence signal (F). The green asterisk indicates a cell expressing mScarlet in which were KHC expression was knocked down; bars = 10 μm. (G–I) Recognition of amyotrophic lateral sclerosis (ALS)-associated KIF5A using recombinant H2. mScarlet-KIF5A(Δexon27) was transfected into CAD cells and stained using recombinant H2; the mScarlet-KIF5A(Δexon27) fluorescence signal (G) and the recombinant H2 fluorescence signal (H). (I) A merged microscopy image showing mScarlet (magenta) and recombinant H2 (green) fluorescence signals; bars = 10 μm.

Next, immunofluorescence microscopy was performed (Fig 2B and C). Recombinant H2 gave cytoplasmic signals in the neuronal CAD cell line (Fig 2B). A similar signal was obtained when the hybridoma-derived H2 mAb was used (Fig 2C). To confirm the specificity of recombinant H2 via immunofluorescence microscopy, we needed a KHC knockdown control cell line. Because H2 recognizes all three KHC isotypes, triple knockdown of KIF5A, KIF5B, and KIF5C was required. To this end, we used a micro (mi)RNA-based knockdown system (Fig 2D). We inserted primary miRNA sequences targeting murine *KIF5A, KIF5B*, and *KIF5C* at the 3’ untranslated region (UTR) of the red fluorescent protein mScarlet. Once expressed, the miRNAs are maturated by the Drosha and Dicer miRNA processing enzymes and are subsequently used by the RNA-induced silencing complex (RISC) to degrade target mRNAs (Cullen, 2004). Therefore, *KIF5A, KIF5B*, and *KIF5C* expression was simultaneously knocked down by the specific miRNAs expressed by the mScarlet-positive CAD cells. 2 days after transfection, the cells were fixed and stained using the fluorescently labelled recombinant H2 (Fig 2E and F). We found that the fluorescence intensity of the H2 signal was significantly weaker in mScarlet-positive cells. These data suggested that the recombinant H2 specifically recognized cytoplasmic KHCs within the CAD cells. Previous studies have used mAbs targeting KHCs, including H1, H2, and SUK4 for immunofluorescence microscopy (Pfister *et al*., 1989; Lippincott-Schwartz *et al*., 1995; Johnson *et al*., 1996). However, these mAbs generate different staining patterns. Our results suggest that H2 is a reliable antibody for use in the immunofluorescence microscopy of mouse cells.

We and others have shown that a specific ALS-associated *KIF5A* mutation induces the aggregation of KIF5A, which leads to its overactivation (Nakano *et al*., 2022; Pant *et al*., 2022). The overactivation of KIF5A, which has been observed both in vitro and in vivo, affects cargo transport. As protein aggregation sometimes masks antibody epitopes, the H2 mAb may not detect ALS-associated KIF5A aggregates efficiently. We therefore wondered whether recombinant H2 could be used to detect ALS-associated KIF5A aggregates in the cell. ALS is caused by mutations that splice out exon 27 of the *KIF5A* gene (Brenner *et al*., 2018; Nicolas *et al*., 2018). The resulting mutant protein, KIF5A(Δexon27), forms aggregates, which are neurotoxic (Nakano *et al*., 2022; Pant *et al*., 2022). We expressed mScarlet-fused KIF5A(Δexon27) in cells and stained them with recombinant H2 mAb. KIF5A(Δexon27) formed aggregates in the cytoplasm as described previously (Fig 2G) (Nakano *et al*., 2022; Pant *et al*., 2022). Immunofluorescence microscopy showed that the recombinant H2 mAb recognized the KIF5A(Δexon27) aggregates in cells (Fig 2H and I). Because antibody detection is more sensitive than the fluorescent protein signal, some aggregates that could not be detected with mScarlet were detected with recombinant H2 (Figs 2G–I). Thus, recombinant H2 is a useful tool for detecting ALS-associated KIF5A aggregates.

### Generation of scFv

scFv is a fusion protein of the variable regions of the IgG heavy and light chains (Trimmer, 2022), which can recognize the same antigen as the original IgG. scFv is encoded by a single cDNA sequence and can therefore be easily manipulated using DNA recombination technology. We synthesized the scFv(H2) using the H2 mAb sequence. The scFv(H2) was fused with the Fc fragment of mouse IgG2 (Fc[mIgG2]). The specificity of the resulting recombinant single-chain antibody, scFv(H2)-Fc(mIgG2), was tested by immunofluorescence microscopy. The scFv(H2)-Fc(mIgG2) could recognize cytoplasmic KHCs in CAD cells (Fig 3B and C). Moreover, this signal was diminished when *KIF5A, KIF5B*, and *KIF5C* were knocked down simultaneously. Thus, the specificity of the recombinant H2 mAb was conserved on its conversion into an scFv.

**Figure 3.**
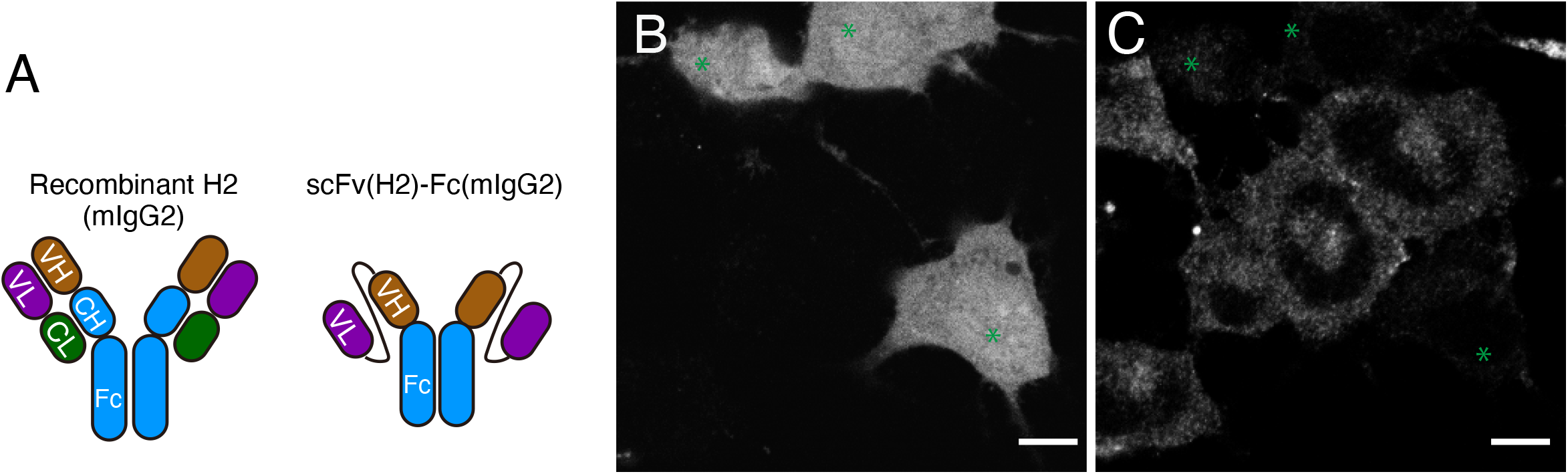
Characterization of the H2-derived scFv. (A) Schematic drawing showing the structures of recombinant H2 and recombinant scFv(H2). (B and C) CAD cells were transfected with the KHC-knockdown vector and stained with the scFv(H2); the mScarlet fluorescence signal (B) and the scFv(H2) fluorescence signal (C). Green asterisks indicate cells expressing mScarlet in which the three KHC isotypes were knocked down; bars =10 μm.

### Diversification by isotype switching

Double and triple staining with different antibodies is a widely used technique to study the function of motor proteins (Kanai *et al*., 2004; Niwa *et al*., 2008). Antibodies generated from different species are generally required for this technique. However, because the mouse is a popular host species used in the production of mAbs, H2 is often difficult to use together with other well-characterized mAbs. To overcome this problem, we fused the scFv(H2) to the Fc domain of the chicken IgY and generated a chickenized scFv(H2) (Fig 4A). By using chickenized scFv(H2), the intracellular localization of KHCs could be visualized using an anti-chicken secondary antibody (Fig 4B and C). To show that the chickenized scFv(H2) did not cross-react with an anti-mouse secondary antibody, chickenized scFv(H2) was used in combination with a mouse anti-tubulin mAb to stain HeLa cells. H2 staining produced a dotted staining pattern in the cytoplasm of HeLa cells, which was similar to the signal observed in CAD cells (Fig 2). Since this staining pattern was not observed in the microtubule channel, we excluded the possibility of cross-reactivity with murine secondary antibodies.

**Figure 4.**
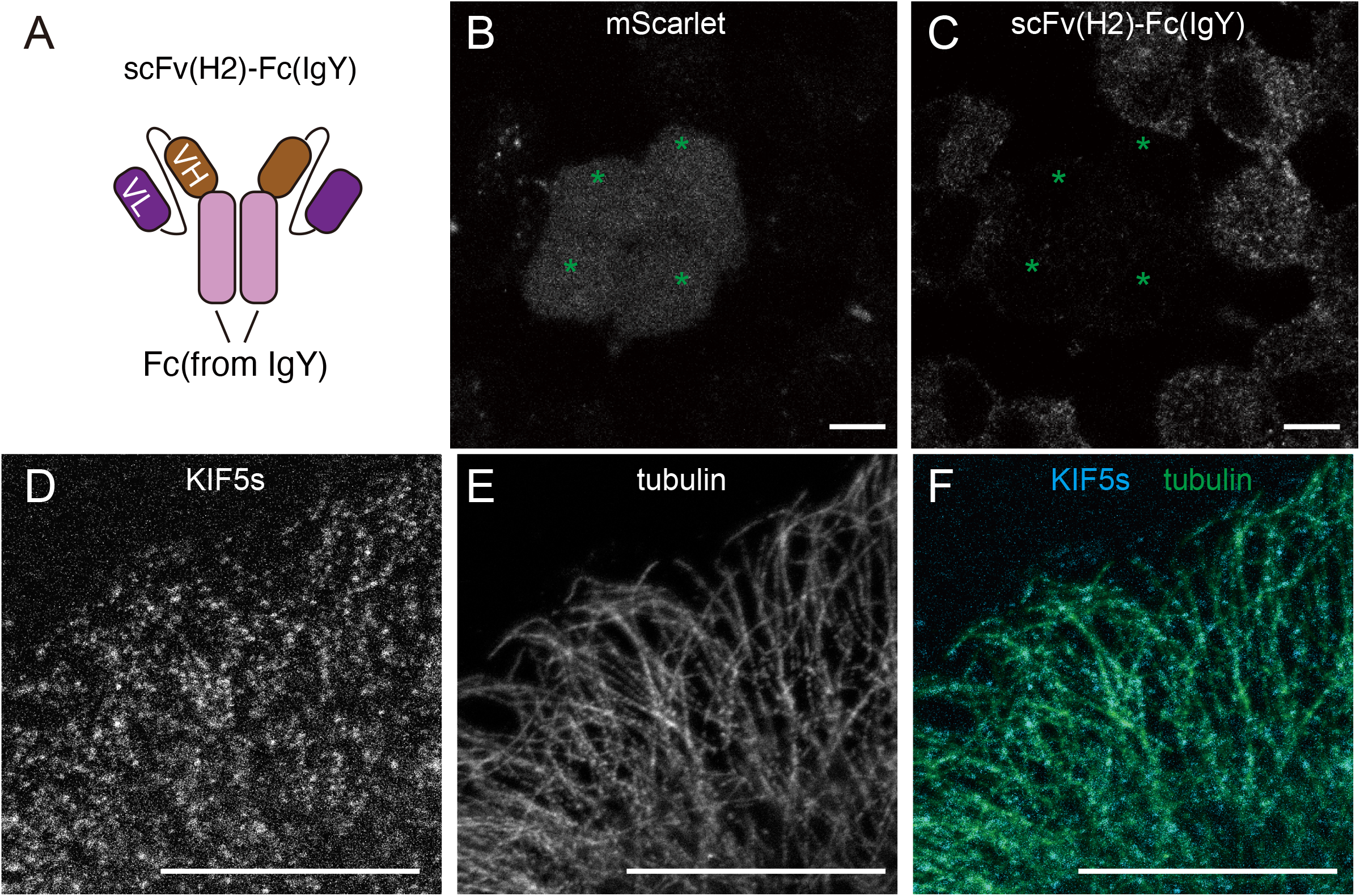
Characterization of the chickenized anti-KHC scFv. (A) A schematic drawing showing the chickenized scFv(H2) structure. (B and C) CAD cells were transfected with the KHC knockdown vector and the localization of the KHCs was visualized using the chickenized scFv(H2); fluorescence signals from mScarlet (B) and the chickenized scFv(H2) (C). Green asterisks indicate CAD cells expressing mScarlet in which the three KHC isotypes were knocked down; bars = 10 μm. (D-F) The KHCs and tubulin were visualized in HeLa cells using the chickenized scFv(H2) and the mouse anti-tubulin antibody, respectively; fluorescence signals from scFv(H2) (D) and tubulin (E). (F) A merged microscopy image of the scFv(H2) (cyan) and anti-tubulin antibody (green) fluorescence signals is shown; bars = 10 μm.

Triple staining was performed to show the advantages of isotype switching (Fig 5). HeLa cells were simultaneously stained using the chickenized scFv(H2), mouse anti-tubulin, and rabbit anti-vimentin antibodies. The experiment demonstrated that KHC colocalized with microtubules and vimentin at the cell periphery. This is consistent with the role of kinesin-1 in driving vimentin filaments along the microtubules (Gyoeva and Gelfand, 1991; Liao and Gundersen, 1998). Moreover, we noticed that both the recombinant H2 mAb and the scFv(H2) produced the same dotted pattern in the cytoplasm (Fig 2 and Fig 5). This may be because kinesin-1 accumulates on cargo vesicles as described previously (Pfister *et al*., 1989). Another possibility is that kinesin-1 forms clusters by liquid-liquid phase separation mechanisms; however, this hypothesis needs further validation. While the original report of H2 staining showed that the KHCs and microtubules did not colocalize (Pfister *et al*., 1989), we found that about 70% of the KHC-positive dots were localized to the microtubules, suggesting that kinesin-1-associated vesicles or kinesin-1 condensates bound to the microtubules.

**Figure 5.**
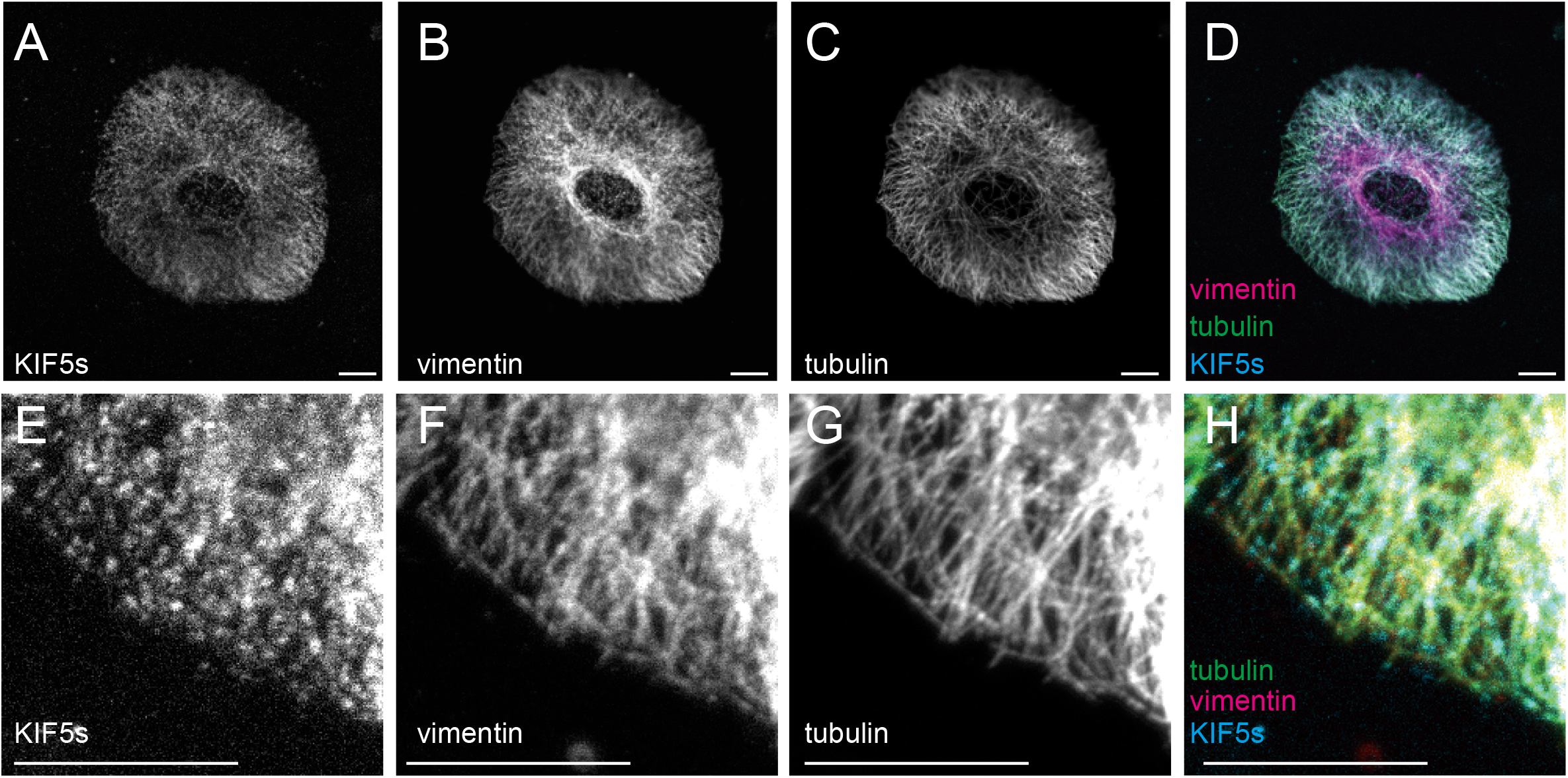
Chickenized anti-KHC scFv enables triple staining. HeLa cells were simultaneously stained with the chickenized scFv(H2), the rabbit anti-vimentin antibody, and the mouse anti-tubulin antibody. Fluorescence signals from the KHC (cyan) (A and E), vimentin (magenta) (B and F), and tubulin (green) (C and G) staining are shown. (D) and (H) show merged images. Bars = 10 μm.

Collectively, our results demonstrate that we successfully generated and diversified the recombinant anti-KHC H2 mAb and the associated scFv(H2). Our experiments have shown how DNA recombinant technology can broaden the utility and application of a classical antibody like H2. While we have not yet succeeded in doing this with the H2 mAb, the scFv can be used as a genetically engineered probe to label endogenous antigens in the cell. This intrabody technique can be used to inhibit the function of antigens in the cell (Trimmer, 2022). However, the scFv is often not soluble in cytosol (Dimitrov *et al*., 2008). Fusion with superfolder GFP (sfGFP) or the ultra-stable cytoplasmic antibody (STAND) could be used to increase the folding and solubility of scFv in the cytoplasm (Tanenbaum *et al*., 2014; Kabayama *et al*., 2020). Future efforts to generate H2 intrabodies will increase the application of recombinant H2 mAbs.

In this report, we have also presented the generation of a triple KIF5 knockdown vector for use in mouse cells. To study the function of kinesin-1, the expression of three KHC isotypes had to be inhibited. While CRISPR-Cas9 can knock out genes in cell lines, using this technology to target three genes simultaneously is time-consuming and laborious. Instead, we initially attempted triple KIF5 knockdown by transfecting mouse cells with three double-stranded RNAs and an mScarlet expression vector. However, we observed no reduction in H2 staining in most of the mScarlet-positive cells (data not shown). By contrast, our triple knockdown plasmid produced highly reproducible results. In 98% of the cells, H2 staining was reduced to the background level. Thus, the triple knockdown plasmid would be a useful tool for studying the function of kinesin-1 in mouse cell lines.

Although antibodies are essential tools in biology, the price of commercially available antibodies is ever increasing, which hampers scientific progress. Thus, the sharing of recombinant antibodies between laboratories will help increase research reproducibility and productivity.

## Materials and Methods

### Cloning of IgG cDNA from H2

Hybridoma lines H1 and H2 were kind gifts from Dr. George Bloom (University of Virginia, VA, USA). Total RNAs from the H1 and H2 hybridomas were purified using the NucleoSpin RNA purification kit (TAKARA, Tokyo, Japan) as described in the manufacturer’s protocol. 5’ RACE was performed using the SMARTer 5’/3’ kit (TAKARA) with slight modifications. Primers for reverse transcription and polymerase chain reaction (PCR) were prepared as previously described (Meyer *et al*., 2019). For reverse transcription, the following primers were used:

5’-TTGTCGTTCACTGCCATCAATC-3’ (for the Kappa chain cDNA)

5’-GGGGTACCATCTACCTTCCAG-3’ (for the Lambda chain cDNA)

5’-AGCTGGGAAGGTGTGCACAC-3’ (for the IgG heavy chain cDNA)

cDNAs linked with a 5’-linker sequence were obtained by reverse transcription, according to the manufacturer’s protocol. Using these cDNAs as templates, PCR reactions were performed using KOD FX neo (Toyobo, Tokyo, Japan), X10 UPM mix primer provided with the SMARTer 5’/3’ RACE kit, and 1 μM of the following gene-specific primers:

5’-<u>GATTACGCCAAGCTT</u> ACATTGATGTCTTTGGGGTAGAAG (for the Kappa chain)

5’-<u>GATTACGCCAAGCTT</u> ATCGTACACACCAGTGTGGC (for the Lambda chain)

5’-<u>GATTACGCCAAGCTT</u> GGGATCCAGAGTTCCAGGTC (for heavy chain)

5’-<u>GATTACGCCAAGCTT</u>-3’ is the sequence used for In-Fusion cloning.

In-Fusion cloning, to insert cDNA into the linearized pRACE vector provided in SMARTer 5’/3’ RACE kit, was performed as described in the manufacturer’s manual. Four independent clones were sequenced by the Sanger method using the Applied Biosystems 3500XL genetic analyzer (Thermo Fisher Scientific, Waltham, MA, USA). As some clones had incorrect mRNA sequences, we selected a sequence that was encoded by at least two independent clones.

### Construction of H2-expressing plasmids

A plasmid expressing an anti-GFP antibody (Andrews *et al*., 2019) was used as a template for recombinant H2 because of its sequence similarity. The variable regions of the anti-GFP heavy and kappa chains in the expression vector were replaced with the equivalent H2 sequences by Gibson assembly, as previously described (Gibson *et al*., 2009). The H2-derived scFv was designed based on the amino acid sequences of the variable regions and synthesized by GeneArt (Thermo Fisher Scientific). The plasmid encoding H2 IgG is available from Addgene; #190690. The plasmid encoding the scFv(H2)-Fc(mouse) is available from Addgene; #190691. The plasmid encoding the scFv(H2)-Fc(chicken) is available from Addgene; #190692.

### Expression of recombinant H2

Recombinant H2 was expressed in Invitrogen™ 293FT cells (Thermo Fisher Scientific). 293FT cells were cultured in Gibco™ DMEM (high glucose) (Thermo Fisher Scientific) supplemented with 10% fetal bovine serum (FBS) (Thermo Fisher Scientific), 500 μg/ml G418 (Nakarai, Kyoto, Japan), 0.1 mM Gibco™ MEM Non-Essential Amino Acids (NEAA) (Thermo Fisher Scientific), and 1 mM Gibco™ MEM Sodium Pyruvate (Thermo Fisher Scientific). DNA transfection was performed as previously described (Longo *et al*., 2013). In brief, 20 μg of plasmid, 100 μl of polyethylenimine (PEI), and 1 ml of Gibco™ OPTI-MEM (Thermo Fisher Scientific) were mixed and incubated for 20 min at room temperature. The mixture was used to transfect 293FT cells in a 75 cm^2^ flask at ∼80% confluency. 48 h after transfection, the culture medium was collected, supplemented with 0.05% NaN_3,_ and stored at 4 °C.

### Preparation of recombinant kinesin

Recombinant kinesin-1 was prepared as previously described (Chiba *et al*., 2022). Human KIF5A, KIF5B, and KIF5C were expressed in Sf9 cells using the Bac-to-Bac Baculovirus Expression system (Thermo Fisher Scientific).

### miRNA plasmids and knockdown

The miRNA sequences were designed as described in the manufacturer’s protocol (Invitrogen). mScarlet-miRNA(KIF5A)-miRNA(KIF5B)-miRNA(KIF5C) was synthesized by GeneArt (Thermo Fisher Scientific). The plasmid is available from Addgene; #190693.

### Knockdown of KHCs and Microscopy

CAD cells were obtained from The European Collection of Authenticated Cell Cultures (ECACC) and cultured in DMEM/F12 (Gibco) supplemented with 8% FBS in a 5% CO_2_ incubator at 37 °C as described previously(Qi *et al*., 1997). For observation, cells were cultured on acid-washed ø18mm coverslips (Matsunami). The miRNA plasmid was transfected using Lipofectamine LTX (Thermo Fisher Scientific) as described in manufacture’s protocol. Two days after transfection, cells were washed with phosphate buffered saline (PBS) and fixed with 4% paraformaldehyde in PBS for two washes. Fixed cells were incubated with the cultured medium obtained from the antibody-expressed HEK293FT cells for 1 hour at 37 °C or overnight at 4 °C. Cells were washed twice with PBS. Mouse anti-tubulin (clone: DM1A, Sigma) and/or rabbit anti-vimentin antibodies (clone: D21H3, Cell Signaling Technology) were diluted in the cultured medium in Fig 4 and 5. Alexa488-labelled anti-mouse IgG antibody (in Fig 2 and 3), Alexa647-labelled anti-chicken IgY and Alexa488-labelled anti mouse IgG antibodies (Fig 4) or Alexa647-labelled anti-chicken IgY, Alexa568-labelled anti-rabbit IgG, and Alexa488-labelled anti-mouse IgG antibodies (in Fig 5) were incubated for 30 min at room temperature. Cells were washed three times with PBS and mounted on the slide glass. An inverted Carl Zeiss Axio Observer Z1 microscope equipped with a 40x/1.3 objective and LSM 800 confocal scanning unit was used as described previously (Chiba *et al*., 2019)

## Acknowledgments

We are very grateful to Dr. George Bloom (University of Virginia, VA, USA) for providing the H1 and H2 hybridomas and allowing us to publish the H2 mAb and scFv sequences and deposit recombinant plasmids to Addgene. SN was supported by JSPS KAKENHI (grants nos. 22H05523 and 20H03247). KC was supported by JSPS KAKENHI (grant no. 21K20621), Uehara Memorial Foundation, and MEXT Leading Initiative for Excellent Researchers (grant no. JPMXS0320200156). We thank Anya Lissina, PhD, from Edanz (https://jp.edanz.com/ac) for editing a draft of this manuscript.

